# Biotinylated surfome profiling identifies potential biomarkers for diagnosis and therapy of *Aspergillus fumigatus* infection

**DOI:** 10.1101/2020.04.03.021493

**Authors:** Lei-Jie Jia, Thomas Krüger, Matthew G. Blango, Olaf Kniemeyer, Axel A. Brakhage

## Abstract

*Aspergillus fumigatus* is one of the most common airborne fungi capable of causing invasive mycoses in immunocompromised patients and allergic diseases in susceptible individuals. In both cases, fungal surface proteins mediate the first contact with the human immune system to evade immune responses or to induce hypersensitivity. Several methods have been established to study the surface proteome (surfome) of *A. fumigatus*, like trypsin shaving, glucanase treatment, or formic acid extraction. Biotinylation coupled with LC-MS/MS identification of peptides is a particularly efficient method to identify the surface exposed regions of proteins that potentially mediate interaction with the host. After biotinylation of surface proteins during spore germination, we detected 314 different surface proteins, including several well-known proteins like RodA, CcpA, and DppV, as well as several allergens, heat shock proteins (HSPs), and previously undescribed surface proteins. Using immunofluorescence microscopy, we confirmed the surface localization of three HSPs, which may have moonlighting functions. Collectively, our study generated a comprehensive data set of the *A. fumigatus* surfome, which complements already existing *A. fumigatus* surface proteome data and allows us to propose a common core set of *A. fumigatus* surface proteins. In addition, our study uncovers the surface-exposed regions of many proteins on the surface of spores or hyphae. These surface exposed regions are candidates for direct interaction with host cells and may represent antigenic epitopes that either induce protective immune responses or mediate immune evasion. Thus, the comprehensive datasets provided and compiled here represent reasonable immunotherapy and diagnostic targets for future investigations.

**HIGHLIGHTS:**

- Surface protein biotinylation coupled with LC-MS/MS analysis provides a comprehensive dataset of the *A. fumigatus* surface proteome.
- 314 different *A. fumigatus* proteins (including immunoreactive proteins, and virulence factors) with surface exposed regions were detected.
- Surface localization of three Hsp70 chaperones was confirmed by protein tagging coupled with immunofluorescence.
- By comparison with other surfome datasets, a core surfome of *A. fumigatus* was defined, which provides possible biomarkers for diagnosis or therapy.

**SIGNIFICANCE:** *Aspergillus fumigatus* is the most important airborne human pathogenic mold, capable of causing both life-threatening invasive pulmonary aspergillosis in immunocompromised patients and allergic infections in atopic individuals. Despite its obvious medical relevance, timely diagnosis and efficient antifungal treatment of *A. fumigatus* infection remains a major challenge. Proteins on the surface of conidia (asexually produced spores) and mycelium directly mediate host-pathogen interaction and also may serve as targets for diagnosis and immunotherapy. However, the similarity of protein sequences between *A. fumigatus* and other organisms, and sometimes even the human host, makes selection of targets for immunological-based studies difficult. Here, using surface protein biotinylation coupled with LC-MS/MS analysis, we identified hundreds of *A. fumigatus* surface proteins with exposed regions, further defining putative targets for possible diagnostic and immunotherapeutic design.

## 1. INTRODUCTION

The saprotrophic fungus *Aspergillus fumigatus*, which occurs on decaying organic material, is associated with a wide spectrum of diseases in humans [1, 2]. The inhalation of *A. fumigatus* airborne conidia may cause life-threatening invasive pulmonary aspergillosis in immunocompromised patients, chronic pulmonary aspergillosis in immunocompetent patients with underlying lung diseases, or allergic infections such as allergic bronchopulmonary aspergillosis (ABPA) in atopic individuals [3, 4]. Despite continuous research and improvements of diagnostic tools, timely diagnosis of *A. fumigatus* remains a challenge [2]. Detection kits for a few different recombinant allergens of *A. fumigatus* are now commercially available for the diagnosis of ABPA, but cross-reactivity with antigens from other microorganisms still makes diagnosis difficult [5]. In addition to DNA and cell wall polysaccharides, fungal proteins exposed to the surface may serve as candidate diagnostic markers and valuable targets for new therapeutics [6, 7].

The *A. fumigatus* cell wall not only maintains cellular integrity and protects from external aggression, but it also serves as a harbor for virulence factors that contribute to immune evasion, adherence, and virulence [8]. Although the cell wall is composed of <10% proteins [2] and hundreds of surface proteins have been detected across various proteome studies [9–11], only a few of these proteins are well characterized, including their roles in *A. fumigatus* virulence. RodA, which forms the hydrophobic rodlet layer on dormant conidia, is the best-studied conidial surface protein. RodA is immunologically inert and can mask dectin-1- and dectin-2-dependent host responses [12, 13]. Despite this important role, a rodletless mutant showed no attenuation in virulence in a mouse infection model of invasive aspergillosis [14]. Another abundant conidial surface protein, CcpA, presumably plays a role in maintaining the spore surface structure and preventing immune recognition. Consequently, it was shown to be essential for virulence in a corticosteroid immunosuppressed mouse infection model [10]. The *A. fumigatus* protein CalA, which is present on swollen conidia and germlings, acts as an invasin through interaction with integrin α_5_β_1_ on host cells and is required for full virulence and lung invasion in corticosteroid-treated mice [15]. Beside these virulence determinants on the conidial surface, many allergens are also surface exposed. Asp f 2 has been described as a zinc acquiring protein and is one of the major allergens of *A. fumigatus* [16]. It was found to bind laminin and IgE antibodies from patients with ABPA [17]. Asp f 2 acts as an immune evasion protein by binding human plasma regulators, which leads to inhibition of opsonization and damage of human lung epithelial cells [18]. However, most of the surface exposed proteins are still uncharacterized, and a comprehensive picture of the *A. fumigatus* surface proteome is lacking, but necessary for a better understanding of the interaction of *A. fumigatus* conidia with the host.

Several proteomic studies have already been performed to define the proteins associated with the cell surface of *A. fumigatus*. These studies relied on either strong acids like formic or hydrofluoric acid (HF) or enzymatic treatments with 1,3-β-glucanase or trypsin to release surface proteins for detection [9, 10, 19]. Around 300 common proteins were detected in formic acid extracts of dormant *A. fumigatus* conidia, while the number significantly increased in *A. fumigatus* mutants lacking either the conidial rodlet-layer or the cell wall polymer, α-1,3-glucan [9]. The combined approach of HF-pyridine extraction and trypsin shaving revealed 477 different proteins on dormant or swollen conidia [10], while 178 different proteins were detected on the surface of *A. fumigatus* conidia during germination using the trypsin shaving approach [11]. In addition, the trypsin shaving approach also showed that the composition of the conidial surface proteome is shaped by the sporulation conditions, such as time and temperature [11].

All the aforementioned methods efficiently extract surface proteins and/or peptides, but they also come with drawbacks: they partially disrupt the surface layer and potentially release cell wall-embedded and cytosolic proteins and peptides in addition to their target cell surface proteins. Cell surface biotinylation has been shown to yield the lowest rate of contamination by cytoplasmic proteins [20] and provides useful information about the surface-exposed protein regions.

Here, we used a widely applied amine-reactive biotinylation method for surface proteomics [15, 21, 22], to label the exposed lysine or N-terminal amino acid residues of *A. fumigatus* cell surface proteins. Subsequently, LC-MS/MS analysis was applied to detect streptavidin-enriched proteins and to define surface-exposed regions based on lysine modifications. The surface localization of several detected proteins was verified through immunofluorescence of Myc-tagged transformants of *A. fumigatus*. Our study complements the picture of the *A. fumigatus* surface proteome and reveals surface proteins that may serve as detection markers or targets for immunotherapy in the future. Perhaps most importantly, we identify a number of proteins that are likely directly involved in the interaction of *A. fumigatus* and the human host.

## 2. MATERIALS AND METHODS

### 2.1 Fungal strains and cultivation

All strains used in this study are listed in Table S1. *A. fumigatus* CEA10 was cultivated as described previously [11]. Briefly, the CEA10 strain was inoculated on *Aspergillus* minimal medium agar plates with 1% (w/v) glucose for 7 days at 37°C. Conidia were harvested in sterile H_2_O and separated from hyphae and conidiophores by filtering (30 μm, MACS^®^). For germination of swollen conidia, 1 × 10^10^ freshly collected conidia were incubated shaking in 50 mL RPMI 1640 (Lonza) for 5 hours (h) at 37°C. 1 × 10^9^ conidia were germinated at the same condition for 8 h to enrich for germlings, and 1 × 10^9^ conidia in 100 mL RPMI 1640 for 14 h to enrich for hyphae.

### 2.2 Biotinylation of surface proteins

Experiments were performed as described previously with minor modification [22]. Conidia, germlings, and hyphae were washed three times with PBS (pH 7.4) and then incubated in 5 mL of PBS containing 5 mg EZ-Link Sulfo-NHS-LC-Biotin (Thermo Fisher Scientific, 21335) for 1 h at 4°C. The reaction was terminated by adding two volumes of 100 mM Tris-HCl, pH 7.4 and incubated further for 30 min. The samples were then washed another three times with PBS (pH 7.4).

### 2.3 Protein extraction and purification

After adding 1 mL of PBS (pH 7.4) containing protease inhibitor (1 tablet per 10 mL, Roche cOmplete^™^, #04693159001) and 500 μL of 0.5 mm diameter glass beads, conidia, germlings, and hyphae were disrupted using a FastPrep homogenizer with the following settings: 6.5 m/s, 3 times for 30 s. The samples were then centrifuged at 16,000 × *g* for 10 min at 4°C. The supernatants were collected and denoted “PBS extracts”. The pellets were washed twice with 1 M NaCl and another three times with 50 mM Tris-HCl, pH 7.4, followed by extraction with SDS buffer (2% (w/v) SDS, 50 mM Tris-HCl, pH 7.4, 100 mM EDTA, 150 mM NaCl and 40 mM DTT) for 10 min at 100°C. For purification of the biotinylated proteins, 1 mg of Roti^®^-Magbeads Streptavidin (Carl Roth, #HP57.1) were equilibrated by three washes in PBS (pH 7.4), then incubated with the protein samples for 2 h in a rotating mixer at 4°C. The beads were washed five times with four different urea buffers (buffer A, 8 M urea, 200 mM NaCl, 2% (w/v) SDS, 100 mM Tris, pH 8; buffer B, 8 M urea, 1.2 M NaCl, 0.2% (w/v) SDS, 100 mM Tris, pH 8, 10% (w/v) ethanol, 10% (w/v) isopropanol; buffer C, 8 M urea, 200 mM NaCl, 0.2% (w/v) SDS, 100 mM Tris, pH 8, 10% (v/v) ethanol, 10% (v/v) isopropanol; buffer D, 8 M urea, 100 mM Tris, pH 8) as previously described [22]. Bound proteins were isolated by incubating the streptavidin beads with 170 μL of elution buffer (30 mM d-biotin, 8 M urea, 2% (w/v) SDS and 100 mM Tris, pH 8) for 30 min at 50°C.

### 2.4 Western blot analysis

Proteins were separated by SDS-PAGE using NuPAGE 4–12% (w/v) Bis-Tris gradient gels (Thermo Fisher Scientific), and then transferred onto a PVDF membrane using the iBlot 2 dry blotting system (Thermo Fisher Scientific). To detect biotinylated proteins, 5 μg of total protein and 20 μL of purified protein were loaded, membranes were blocked with Western Blocking Reagent (Roche), and then incubated with Pierce^®^ streptavidin-HRP (1:2,000; Thermo Fisher Scientific, #21130) overnight at 4°C. To detect the Myc-tagged proteins, 10 μg of total protein were loaded, membranes were blocked with a 5% (w/v) solution of skim milk, and then incubated with primary antibody (1:1,000; Myc-tag mouse mAb, Cell Signaling Technology, #2276) overnight at 4°C. Hybridization with a secondary antibody (HRP-linked anti-mouse IgG, Cell Signaling Technology, #7076) was performed for 1 h at room temperature. Chemiluminescence of HRP substrate was detected with the Fusion FX7 system (Vilber Lourmat, Germany).

### 2.5 Immunofluorescence microscopy

The fungal materials were washed three times with PBS (pH 7.4) and then blocked with 1% (w/v) bovine serum albumin (BSA) in PBS for 1h at room temperature. For the detection of surface biotinylation, *A. fumigatus* conidia, germlings, and hyphae were incubated with Alexa Fluor^™^ 635 conjugated streptavidin (1:100; Thermo Fisher Scientific, S32364) for 1 h in the dark. The surface localization of Myc-tagged proteins, was examined using an anti-Myc primary antibody (1:100 of Myc-tag mouse mAb, Cell Signaling Technology, #2276) for 2 h at room temperature and a secondary antibody (1:500; Donkey anti-mouse IgG H&L, Alexa Fluor^®^ 568, Abcam #ab175472) for 1 h at room temperature in the dark. After three times washing with PBS, the samples were examined under a Zeiss Axio Imager M2 microscope.

### 2.6 In-solution protein digest with trypsin

150 μL of eluted protein samples were reduced by adding 4 μL of 500 mM TCEP (tris(2-carboxyethyl)phosphine in 100 mM TEAB) for 1 h at 55°C. 4 μL 625 mM iodoacetamide was added to each sample and incubated for 30 min at room temperature in the dark. Proteins were then precipitated using the methanol-chloroform-water method [23]. Protein samples were resolubilized in 100 μL of 100 mM triethylammonium bicarbonate (TEAB) and sonicated for 10 min. Protein content was measured with a Merck Millipore Direct Detect infrared spectrometer. The samples were treated with trypsin/Lys-C protease mix (Promega, V5072) at a protease:protein ratio of 1:25 for 16 h at 37°C. The reaction was stopped with 10 μL of 10% (v/v) formic acid and evaporated using a SpeedVac (Thermo Fisher Scientific). Peptides were resolubilized in 25 μL 0.05% (v/v) trifluoroacetic acid (TFA) and 2% (v/v) acetonitrile (ACN) in water and sonicated for 15 min in a water bath before transfer to a 10 kDa molecular weight cut-off filter. After 15 min of centrifugation at 14,000 × g (4°C), the samples were then transferred to HPLC vials and stored at −20°C.

### 2.7 Genetic manipulation of *A. fumigatus*

All oligonucleotides utilized in this study are listed in Table S2. For surface localization studies, a pLJ-*Ssc70-Myc* construct was generated using pTH1 plasmid [24] as backbone. A 3,276 bp DNA fragment containing 1,010 bp 5’ promoter region, the *ssc70* gene without TAA and an in-frame fused Myc-tag coding sequence was PCR amplified from genomic DNA using primers Ssc70-M_F and Ssc70-M_R. Another 371 bp DNA fragment was amplified from pTH1 plasmid using primers Myc_F and pTH_R. These two fragments were mixed as the template and amplified using primers Ssc70-M_F and pTH_R to generate the *Ssc70-Myc* fragment. *Ssc70-Myc* fragment was digested with *Kpn*I and *Not*I, and then inserted to pTH1 plasmid using T4 ligase (Thermo Fisher Scientific) to yield the pLJ-*Ssc70-Myc* plasmid. The pLJ-*Ssc70-Myc* plasmid was then used as the backbone to generate other Myc-tag constructs. For example, a 3,073 bp DNA fragment containing *hsp70* gene and the promoter region was PCR amplified from genomic DNA using primers Hsp70-M_F and Hsp70-M_R. The DNA fragment was then inserted into the *Kpn*I and *Hin*dIII digested pLJ-*Ssc70-Myc* backbone using the CloneEZ^™^ PCR Cloning Kit (GenScript) to get pLJ-*Hsp70-Myc* plasmid. Primers BipA-M_F and BipA-M_R were used to amplify the *bipA* gene with promoter region. Primers Ssz-M_F and Ssz-M_R were used to amplify the *ssz* gene with promoter region. The plasmids were used to transform protoplasts from *A. fumigatus* strain A1160 (CEA17 Δ*akuB*^KU80^) [25].

### 2.8 LC-MS/MS analysis

LC-MS/MS analysis of tryptic peptides was performed on an Ultimate 3000 RSLC nano instrument coupled to a QExactive HF mass spectrometer (both Thermo Fisher Scientific). Tryptic peptides were trapped for 4 min on an Acclaim Pep Map 100 column (2 cm × 75 μm, 3 μm) at a flow-rate of 5 μL/min. The peptides were then separated on an Acclaim Pep Map column (50 cm × 75 μm, 2 μm) using a binary gradient (A: 0.1% (v/v) formic acid in H_2_O; B: 0.1% (v/v) formic acid in 90:10 (v/v) ACN/H_2_O): 0 min at 4% B, 6 min at 8% B, 30 min at 12% B, 75 min at 30% B, 85 min at 50% B, 90–95 min at 96% B, 95.1–120 min at 4 % B. Positively charged ions were generated by a Nanospray Flex Ion Source (Thermo Fisher Scientific) using a stainless steel emitter with 2.2 kV spray voltage. Ions were measured in data-dependent MS2 (Top15) mode: Precursor ions (z=2–5) were scanned at m/z 300–1,500 (resolution, R: 120,000 FWHM; automatic gain control, AGC target: 3·10^6^; maximum injection time IT_max_: 120 ms). Fragment ions generated in the HCD cell at 30% normalized collision energy using N_2_ were scanned (R: 15,000 FWHM; AGC target: 2·10^5^; max. IT: 90 ms) using a dynamic exclusion of 30 s.

### 2.9 Database search and data analysis of trypsin-cleaved surface peptides

The MS/MS data were searched against the Aspergillus Genome database (AspGD) of *Aspergillus fumigatus* Af293 (http://www.aspergillusgenome.org/download/sequence/A_fumigatus_Af293/current/;2019/02/03 YYYY/MM/DD) using Proteome Discoverer (PD) 2.2 and the algorithms of Mascot 2.4.1, Sequest HT, and MS Amanda 2.0. Two missed cleavages were allowed for tryptic peptides. The precursor mass tolerance was set to 10 ppm and the fragment mass tolerance was set to 0.02 Da. Dynamic modifications were set as oxidation (+15.995 Da) of Met and NHS-LC-Biotin (+339.162 Da) modification of Lys and the protein N-terminus. The static modification was set to carbamidomethylation (+57.021 Da) of Cys. One unique rank 1 peptide with a strict target false discovery (FDR) rate of < 1% on both peptide and protein level (compared against a reverse decoy database) were required for positive protein hits. Protein abundance quantification was performed by the Minora algorithm of PD2.2 (area under the curve approach).

### 2.10 Data availability

The mass spectrometry proteomics data have been deposited to the ProteomeXchange Consortium *via* the PRIDE [26] partner repository with the dataset identifier PXD018071.

## 3. RESULTS

### 3.1 Biotinylation of *A. fumigatus* surface proteins

In this study, we sought to expand the repertoire of *A. fumigatus* surface proteins and their surface-exposed epitopes, which could directly mediate pathogen-host interaction and serve as targets for diagnosis and therapy. We used the biotinylation reagent sulfosuccinimidyl-6-(biotinamido)hexanoate (Sulfo-NHS-LC-Biotin) to label the surface-exposed primary amine residues (*e.g*., the ε-amino group of lysine residues) throughout germination of *A. fumigatus* conidia (Figure 1). The water-soluble and negatively charged Sulfo-NHS-LC-Biotin is one of the most frequently applied biotinylation reagents, particularly in regard to surface proteomics [21, 22, 27]. Samples were collected from dormant conidia grown at 37°C on AMM agar plates, swollen conidia (5 h), germlings (8 h), and hyphae (14 h) germinated at 37°C in RPMI liquid medium as described previously [11]. Although there is a possibility of biotin reagent permeation into the fungal cells [22], the biotinylation signal was confined mainly to the surface of conidia and hyphae as observed by immunofluorescence staining against biotinylated proteins (Figure 2A and Figure S1). Due to the high rigidity of the *A. fumigatus* cell wall, the fungal samples were first disrupted by glass beads in PBS buffer to release loosely attached surface proteins. This was followed by a second step using SDS buffer extraction to remove non-covalently bound hydrophobic surface proteins (Figure 1). After purification, the samples were checked by Western blot for the level of protein biotinylation and recovery from the streptavidin beads (Figure 2B and Figure S2). Proteins, varying widely in molecular mass were detected, demonstrating the ability to label a wide range of proteins using this approach. We also observed biotinylated protein bands in non-biotinylated samples (Figure 2B and Figure S2). Most likely, these represent the three known biotin-dependent carboxylases in *A. fumigatus*, encoded by Afu4g07710, Afu2g08670, and Afu5g08910, which each showed high abundance according to the observed peptide spectrum matches (PSM) in our LC-MS/MS analyses (Dataset 1). They are easily distinguishable from the NHS-LC-Biotin modification, which adds 339 dalton of mass to the modified proteins.

**Figure 1.**
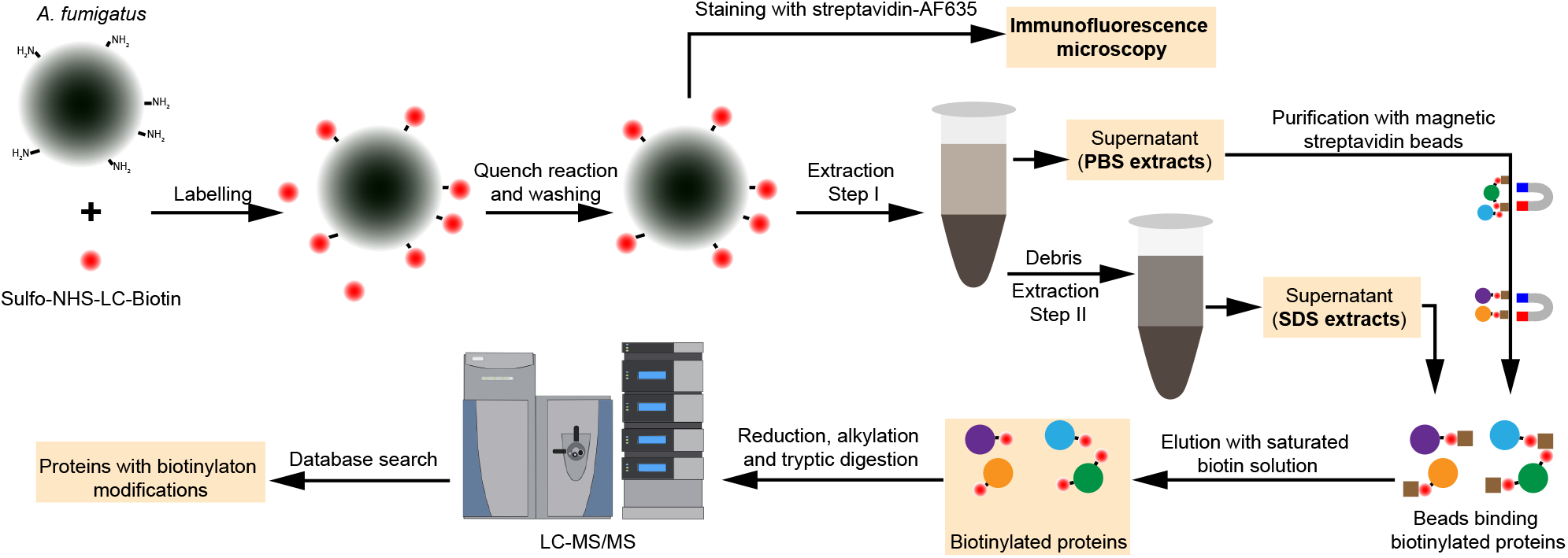
Flowchart for the biotinylation and purification of *A. fumigatus* surface proteins. The procedure starts with covalent labeling of surface proteins with Sulfo-NHS-LC-Biotin for 30 min at 4°C. The fungal material was broken with glass beads in PBS buffer to release the loosely attached surface proteins (PBS extracts). Some non-covalently linked cell wall proteins were extracted using SDS buffer. Biotinylated proteins were purified using magnetic streptavidin beads and then analyzed by LC-MS/MS.

**Figure 2.**
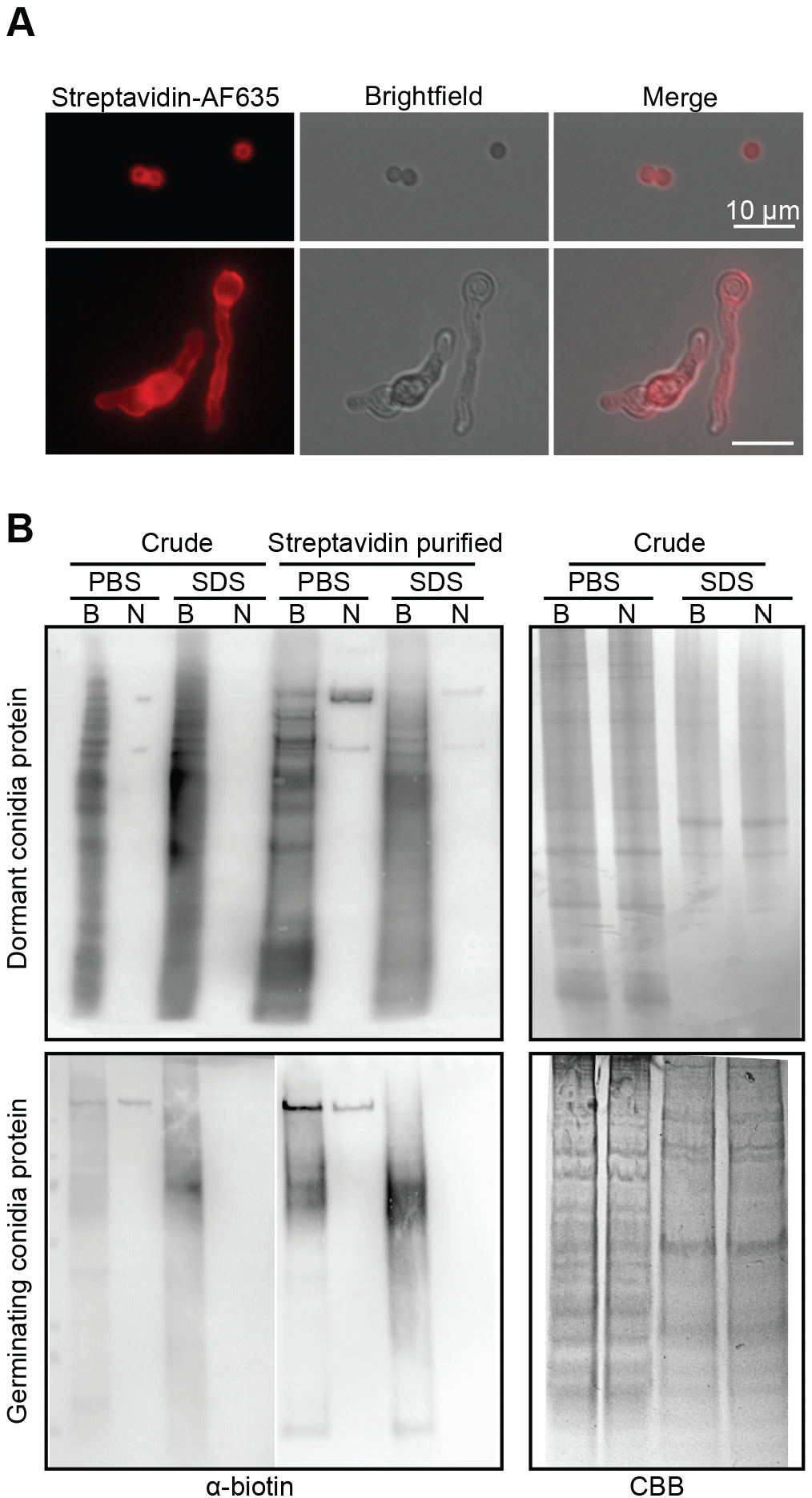
Detection of *A. fumigatus* surface protein biotinylation. (A) Immunofluorescence staining of *A. fumigatus* dormant conidia and germinating conidia with Alexa Fluor^™^ 635 conjugated streptavidin (Streptavidin-AF635). Scale bar is 10 μm. (B) Immunoblotting analysis of the crude protein extracts and purified proteins from dormant and germinating conidia. B, biotinylated; N, non-biotinylated; PBS, PBS extracts; SDS, SDS buffer extracts; CBB, Coomassie brilliant blue.

### 3.2 Identification of constitutively, and stage-specifically, exposed proteins

LC-MS/MS analysis resulted in a proteome set consisting of 1139 different *A. fumigatus* proteins, which were detected with at least 2 peptides and/or a PSM value ≥ 10. Approximately 28% (314) of the detected proteins had a detectable NHS-LC-Biotin modification when considering all germination conditions (Figure 3A). 77 proteins had a predicted signal peptide and 22 proteins had at least one transmembrane domain (fungidb.org; [28]). Over the course of germination, 74 proteins with biotinylation were identified on dormant conidia, 75 on swollen conidia, 93 on germlings, and 214 on hyphae. Most of the proteins detected with biotin modifications were found in PBS extracts, only a few (5 to 14) were exclusively found in SDS extracts (Figure 3A, 3B, and Dataset 1). In accordance with our previous surface proteomics study based on a trypsin-shaving approach [11], our data confirm the dynamic change of the surface-exposed proteome of *A. fumigatus* across germination. There were 24, 16, 30, and 146 proteins detected exclusively on dormant, swollen, germinating conidia, and hyphae, respectively (Figure 3C).

**Figure 3.**
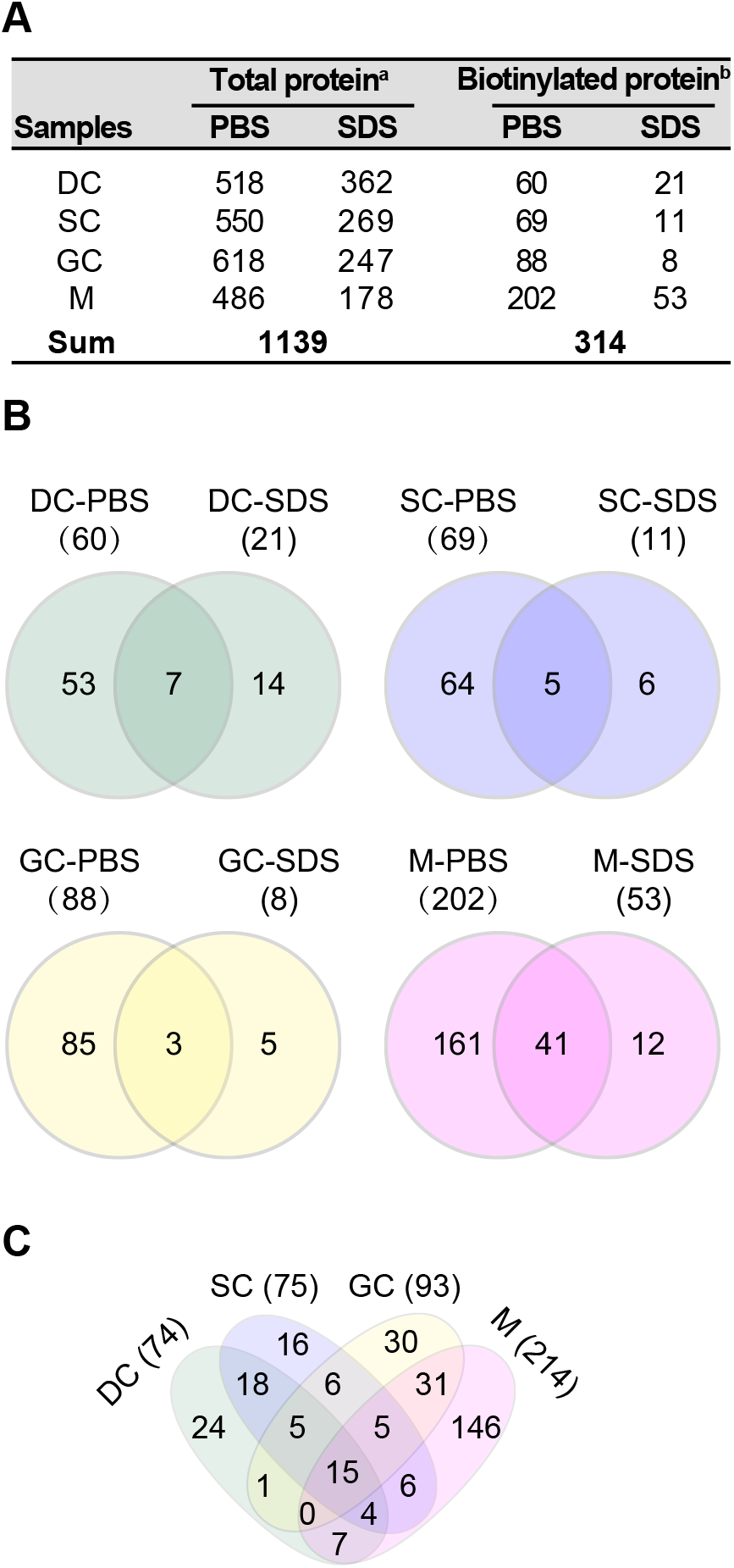
Overview of the surface proteome identified by biotinylation coupled with LC-MS/MS analysis. (A) Number of proteins identified in different samples. ^a^, number of proteins identified by at least two unique peptides or with PSMs (peptide spectrum matches) ≥ 10 were considered. ^b^, number of proteins detected with at least one peptide with biotinylation modification. (B and C) Venn diagrams showing the common and specific surface proteins with biotinylation across different extractions (B) and developmental stages (C).

Throughout germination, 15 proteins were biotinylated in all four stages (Table 1), including several already described surface proteins, for example, RodA [12], the translational elongation factor 1 alpha Tef1 [29], the peptidase DppV [30], the 1,3-beta-glucanosyltransferase Bgt1[31] and Bgt2 [32], the 14-3-3 protein ArtA [29], and the MedA-regulated, putative adhesin Afu3g00880 [33]. The biotinylation of RodA K50 was detected in all four stages, while the biotinylation of K55 and K126 was only detected in dormant and swollen conidia (Table 1). All lysine residues are localized in α-helical regions of the protein [9]. RodA also exhibited high abundance in conidia but not mycelium (Table 2 for top 15). Tef1 was detected mainly in SDS extracts (Table S3). The biotinylation of Tef1 was detected at sites K472, K476, K483, and K486, indicating that the C-terminus of Tef1 was exposed on the conidial and mycelial surface. Somewhat surprisingly, the histone H2B (Htb1, Afu3g05350) seemed to be one of the most abundant proteins throughout the germination course (Table 1 and 2). The total PSM value of Htb1 in dormant conidia was even higher than that of RodA (Dataset 1) in contrast to previous studies using different approaches [10]. One possible explanation for this difference is the difference in the total number of the modifiable lysines in each protein. There are 23 lysine residues in the 140 amino acid residues (AAs) of Htb1, 9 of which were detected with a biotinylation modification (Table 1). In contrast, there are only 9 lysine residues present in the hydrophobic protein RodA.

**Table 1.**
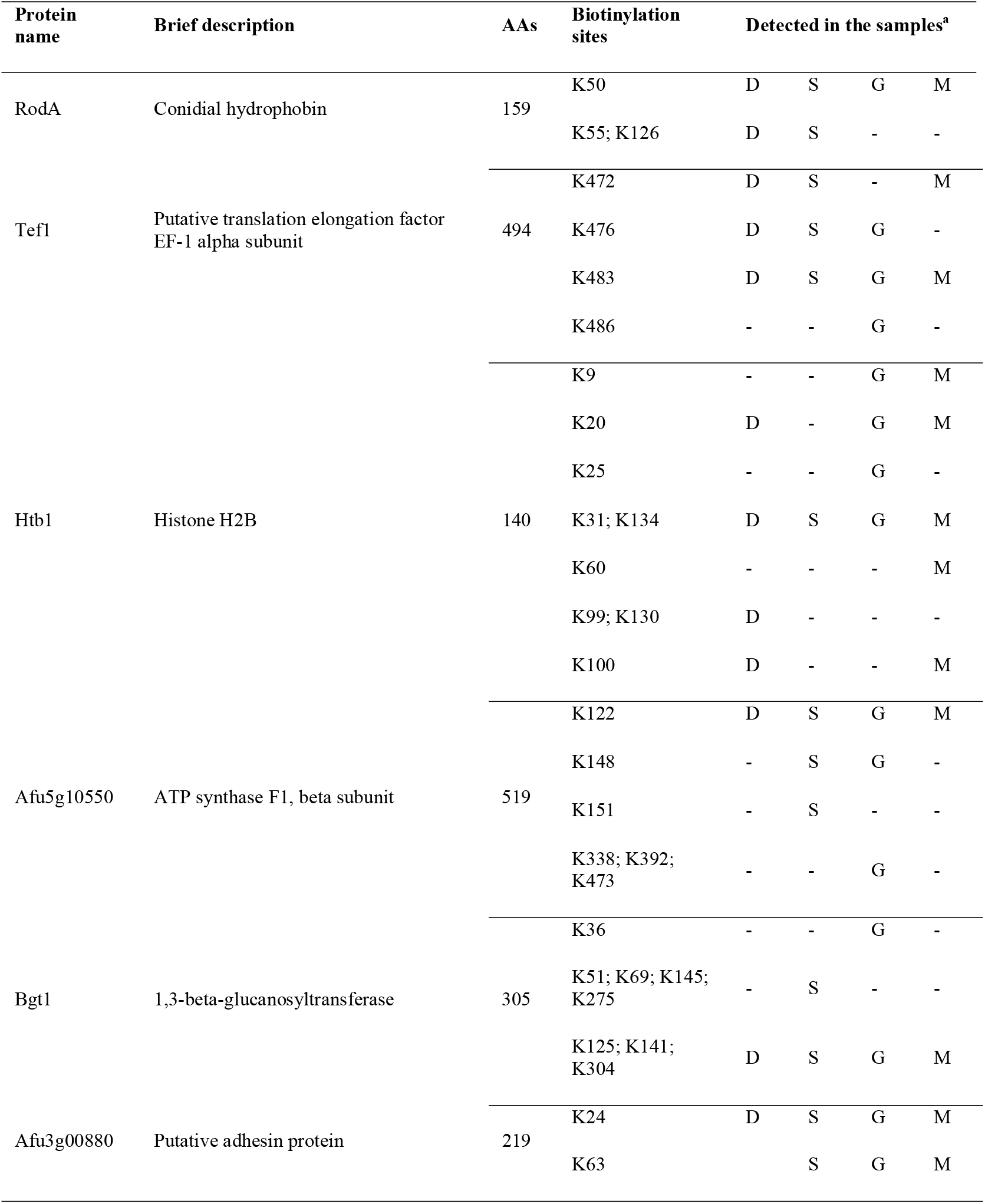
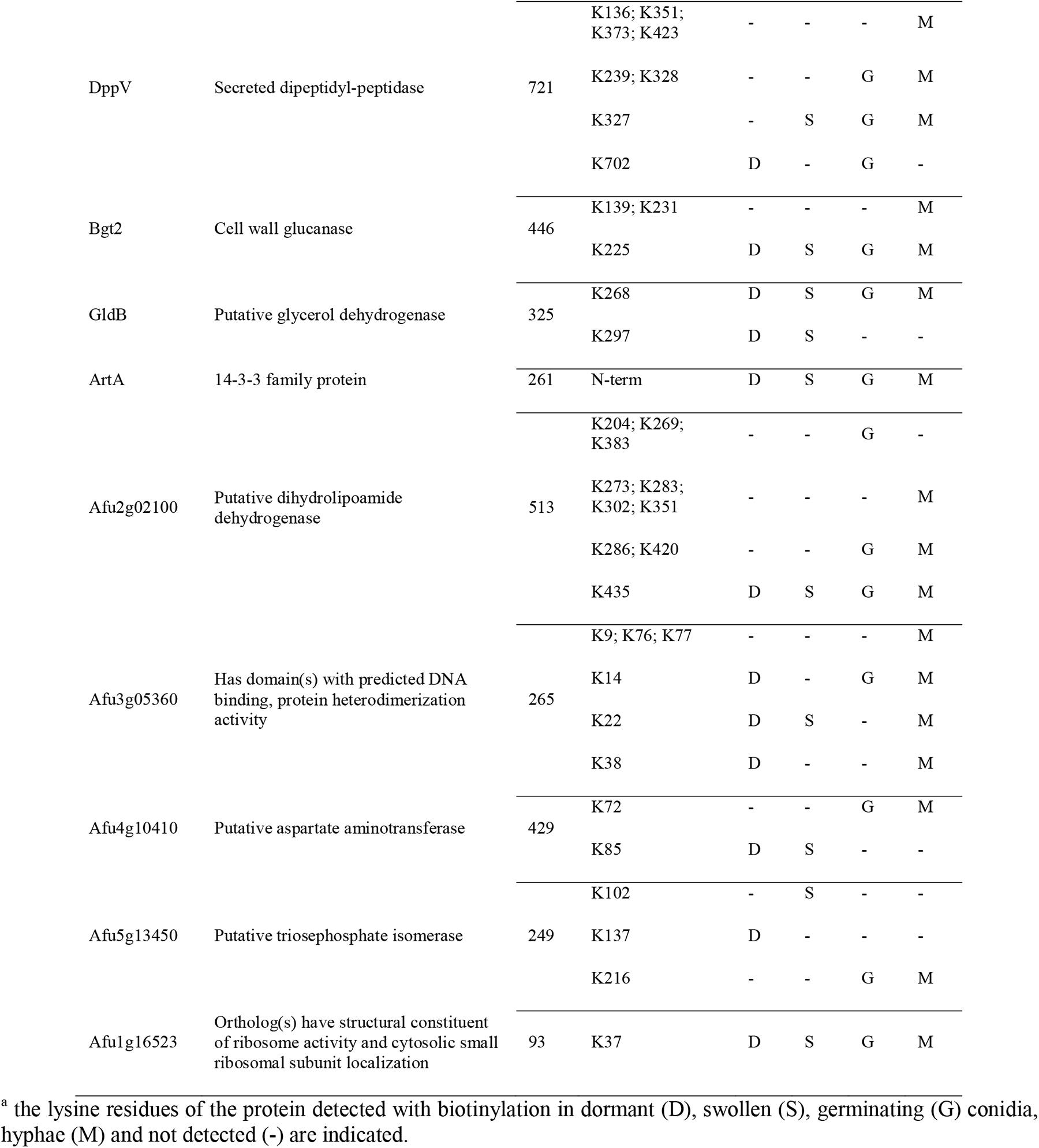
Proteins identified in all stages throughout germination in the streptavidin-enriched fractions.

**Table 2.**
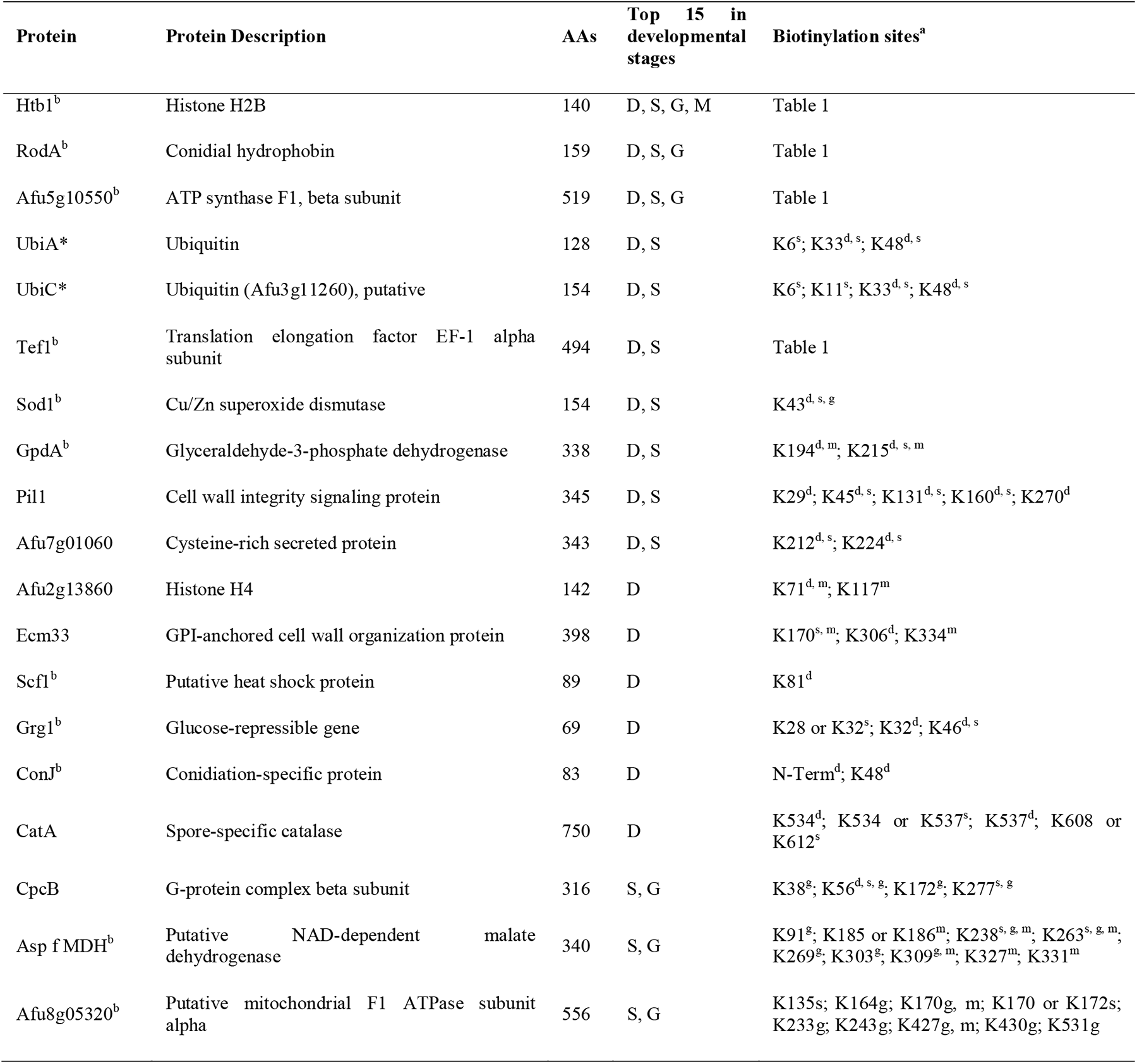
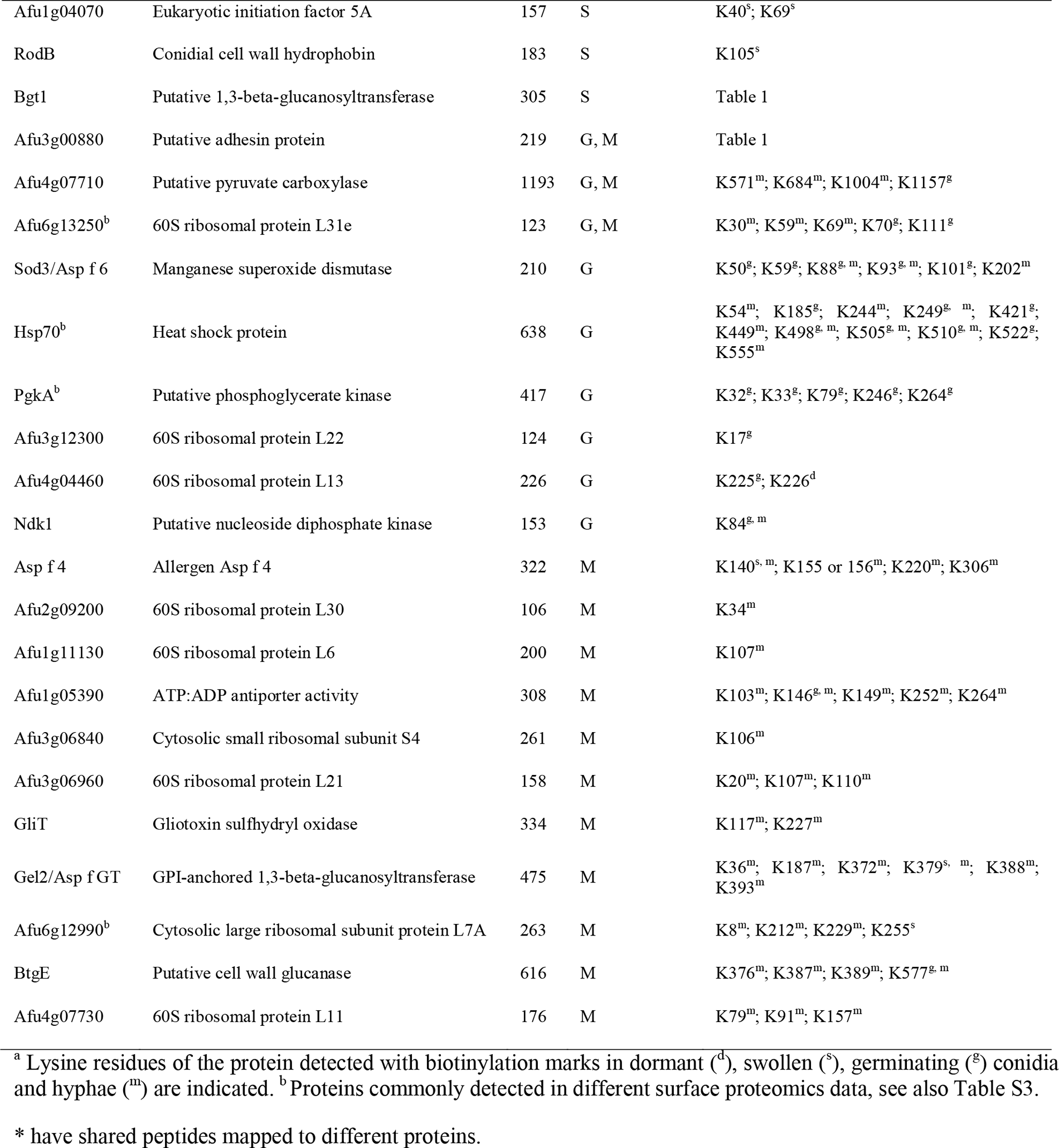
LC-MS/MS analysis of highly abundant proteins detected throughout the germination course (top 15 of each morphotype).

### 3.3 Immunoreactive proteins are exposed on the surface of *A. fumigatus*

In a unique subset of atopic individuals, sensitization to *A. fumigatus* allergens can develop into allergic asthma, allergic sinusitis, and following fungal lung colonization into ABPA [2]. Twenty-three different allergens have been reported in accordance to the systematic allergen nomenclature (www.allergome.org), but actually more than 100 immunoreactive *A. fumigatus* proteins have already been detected by immunoproteomic studies [5]. In our work, 22 allergens (Table 3) were detected by surface biotinylation. The allergens Asp f 17, Asp f 18, Asp f 27, Asp f Mannosidase, Asp f Catalase, and Asp f Glucosidase were detected on dormant conidia. The biotinylation of Asp f 27 K140 was detected on dormant, swollen, and germinating conidia. In addition to Asp f 27, three allergens (Asp f 9, Asp f FDH, and Asp f MDH) were detected on swollen conidia, germlings, and hyphae. Biotinylation of Asp f 9 K190, Asp f MDH K238, and K263 were detected on the three different morphologies of *A. fumigatus*. Ten allergens were only detected on germlings and/or hyphae (Table 3). Consistent with the literature, these data again clearly demonstrate that numerous known *A. fumigatus* allergens are exposed on the fungal surface [9–11, 18, 34].

**Table 3.** *A. fumigatus* allergens exposed to the surface.

| Allergen<sup>a</sup> | Protein Description | AAs | Detected<br>in the<br>samples | Biotinylation sites |
| --- | --- | --- | --- | --- |
| Asp f 1 | Mitogillin | 176 | G, M | K96<sup>g, m</sup>; K138<sup>g</sup>; K140<sup>g</sup> |
| Asp f 2 | Allergen Asp f 2 | 314 | M | K83<sup>m</sup> |
| Asp f 4 | Allergen Asp f 4 | 322 | S, M | K140<sup>s, m</sup>; K155 or 156<sup>m</sup>; K220<sup>m</sup>; K306<sup>m</sup> |
| Asp f 6 (Sod3) | Putative manganese superoxide dismutase | 210 | G, M | K50<sup>g</sup>; K59<sup>g</sup>; K88<sup>g, m</sup>; K93<sup>g, m</sup>; K101<sup>g</sup>; K202<sup>m</sup> |
| Asp f 9 | Cell wall glucanase | 395 | S, M, G | K190<sup>s, g, m</sup> |
| Asp f 11 (Cyp4) | Putative cyclophilin | 205 | S | K119<sup>s</sup> |
| Asp f 12 (Hsp90) | Heat shock protein | 706 | G | K481<sup>g</sup> |
| Asp f 17 (Mp1)<sup>b</sup> | Putative cell wall galactomannoprotein | 284 | D | K63<sup>d</sup>; K144<sup>d</sup> |
| Asp f 18 (Alp2) | Autophagic (vacuolar) serine protease | 495 | D | K261<sup>d</sup>; K268<sup>d</sup>; K271<sup>d</sup>; K291<sup>d</sup> |
| Asp f 23 (RpL3) | Allergenic ribosomal L3 protein | 392 | M | K5<sup>m</sup> |
| Asp f 27<sup>b</sup> | Putative peptidyl-prolyl cis-trans isomerase | 163 | D, S, G | K43<sup>d, s</sup>; K140<sup>d, s, g</sup>; K152 or K153<sup>d</sup> |
| Asp f 28<sup>b</sup> | Putative thioredoxin | 171 | M | K125<sup>m</sup> |
| Asp f Mannosidase (MsdS)<sup>b</sup> | Putative 1,2-alpha-mannosidase | 503 | D | K212<sup>d</sup>; K274<sup>d</sup>; K415<sup>d</sup>; K416<sup>d</sup>; K488<sup>d</sup> |
| Asp f Catalase (Cat1)<sup>b</sup> | Catalase | 728 | D | K261<sup>d</sup>; K346<sup>d</sup>; K466<sup>d</sup> |
| Asp f Glucosidase (Exg12)<sup>b</sup> | Secreted beta-glucosidase | 863 | D, M | K127<sup>d</sup>; K178<sup>d</sup>; K415<sup>d</sup>; K603<sup>d</sup>; K839<sup>d</sup>; K843<sup>d, m</sup> |
| Asp f FDH | Putative NAD-dependent formate dehydrogenase | 418 | S, M, G | K304<sup>m</sup>; K319<sup>s, g</sup>; K327<sup>m</sup> |
| Asp f MDH<sup>b</sup> | Putative NAD-dependent malate dehydrogenase | 340 | S, M, G | K91<sup>g</sup>; K185 or K186<sup>m</sup>; K238<sup>s, g, m</sup>; K263<sup>s, g, m</sup>; K269<sup>g</sup>; K303<sup>g</sup>; K309<sup>g, m</sup>; K327<sup>m</sup>; K331<sup>m</sup> |
| Asp f GT (Gel2) | GPI-anchored 1,3-beta-glucanotransferase | 475 | S, M | K36<sup>m</sup>; K187<sup>m</sup>; K372<sup>m</sup>; K379<sup>s, m</sup>; K388<sup>m</sup>; K393<sup>m</sup> |
| Asp f GPI | Putative glucose-6-phosphate isomerase | 553 | M | K35<sup>m</sup>; K451<sup>m</sup> |
| Asp f AT_V | Putative class V aminotransferase | 386 | M | K381<sup>m</sup> |
| Asp f gamma_Actin (Act1)<sup>b</sup> | Actin | 393 | M | K346<sup>m</sup> |
| Asp f RPS3<sup>b</sup> | 40S ribosomal protein S3 | 266 | M | K78<sup>m</sup> |

<sup>a</sup> *A. fumigatus* allergens based on allergome website (www.allergome.org). <sup>b</sup> Allergens commonly detected in different surface proteomics data, see also Table S3.

In addition to the allergens, there were also other immunoreactive proteins that could serve as biomarkers for diagnosis or targets for allergen immunotherapy [35–37]. DppV, Bgt1, Bgt2, and Afu5g10550 were detected with biotinylation modifications in all four morphotypes (Table 1). Afu5g10550 encodes an ATP synthase F1 beta subunit, which reacts with immunosera from rabbits exposed to *A. fumigatus* conidia [35]. Biotinylation of Afu5g10550 K122 was detected throughout germination. It was also one of the most prevalent proteins detected on dormant, swollen, and germinating conidia (Table 1 and 2). The immunoreactive GpdA, Asp f MDH, Bgt1, Asp f 6, PgkA, Asp f 4, GliT, Asp f GT, and Hsp70 were also prevalent on the surface of one or two morphological stages (Table 2).

### 3.4 Heat shock proteins are exposed to the surface

Heat shock proteins (HSPs) and a large set of co-chaperones are ubiquitous molecular chaperones that act in maintaining protein homeostasis [38]. It has been well established that HSPs are also secreted extracellularly or localized on the cell surface [39]. Hsp90, also known as Asp f 12, is detectable on the cell wall of *A. fumigatus* [34] and it was also found to be biotinylated in this study (Table 3). Considering the detection of biotinylated Hsp90 and Hsp70, we investigated whether other chaperone-related proteins are present on the cell surface of *A. fumigatus*. Indeed, at least 16 chaperone-related proteins were detected with biotinylation modifications (Table 4). Most of the chaperones were detected on the surface of germlings or hyphae with the exception of Scf1 and GrpE. Scf1 shows similarities to the 12 kDa heat shock protein of *S. cerevisiae* and was biotinylated at K81 (Table 4). It is one of the most prevalent proteins on dormant conidia of the *A. fumigatus* strains CEA10 (Table 2) and ATCC 46645 [10]. Six chaperones (Hsp70, Hsp88, HscA, BipA, Ssc70, and Lhs1) belonging to the Hsp70 family were detected in our study (Table 4). In addition to Hsp70, four other Hsp70 family proteins (Sti1, Hsp88, HscA, and Ssc70) have shown to induce serological antibody responses in ABPA patients [37].

**Table 4.**
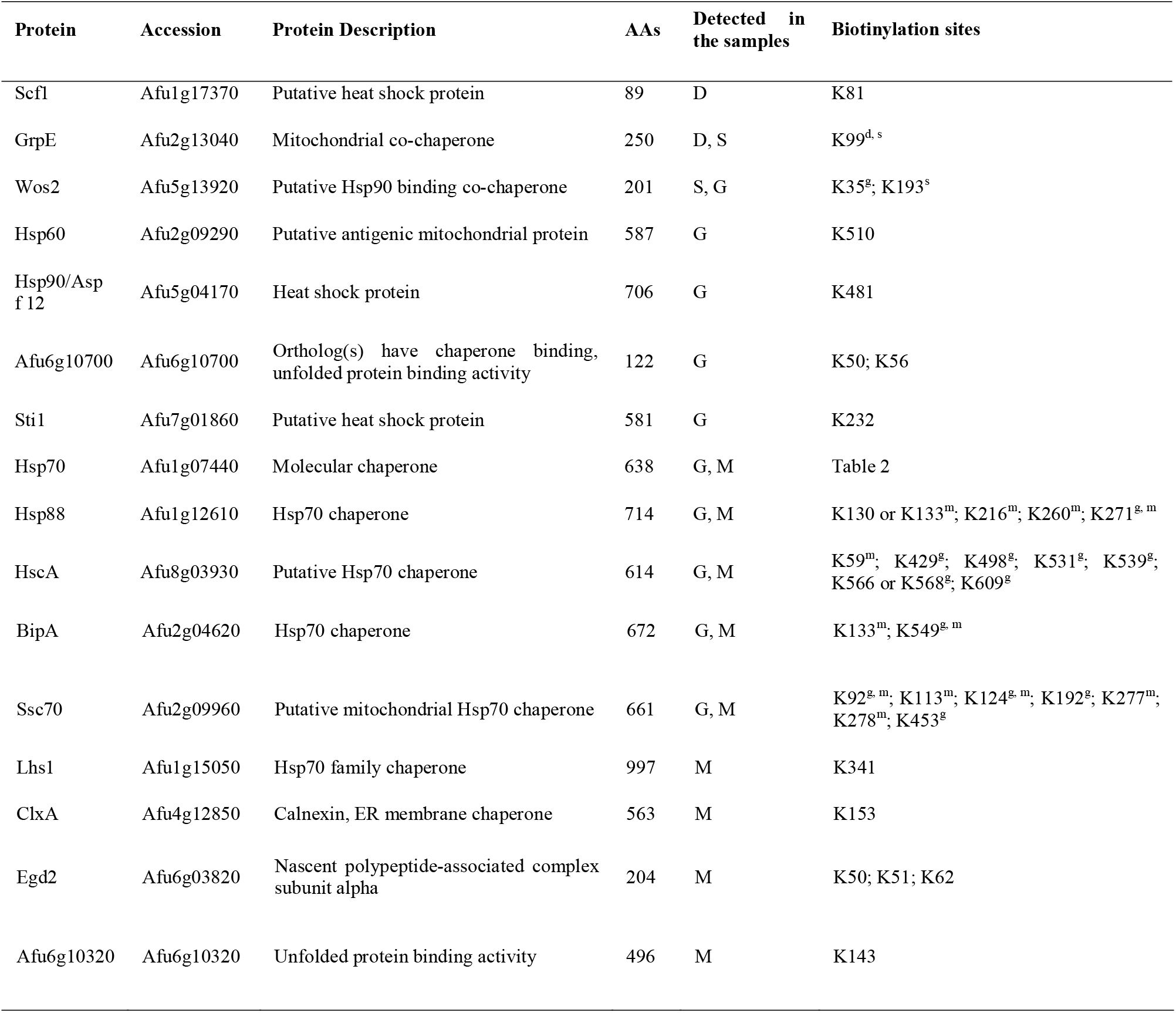
Chaperones and co-chaperones with cell surface localization.

To confirm the surface localization of the HSPs, we generated constructs by fusing a Myc-tag to the C-terminus of the HSPs. These constructs were ectopically integrated into the genome of *A. fumigatus* (Supplementary Table 1 and 2). The *hsp70-Myc, ssc70-Myc, bipA-Myc* and *ssz-Myc* transformants were verified by immunoblotting using an anti-Myc tag antibody (Figure S3). Using immunofluorescence microscopy, we found that all of the Myc-tagged fusion proteins were localized on the surface (Figure 4), except for the negative control, the HSP 70 chaperone Ssz (Afu2g02320), which was not detected on the surface in our biotinylation experiment. This independently confirms that HSPs of *A. fumigatus* are localized to the cell surface of the fungus.

**Figure 4.**
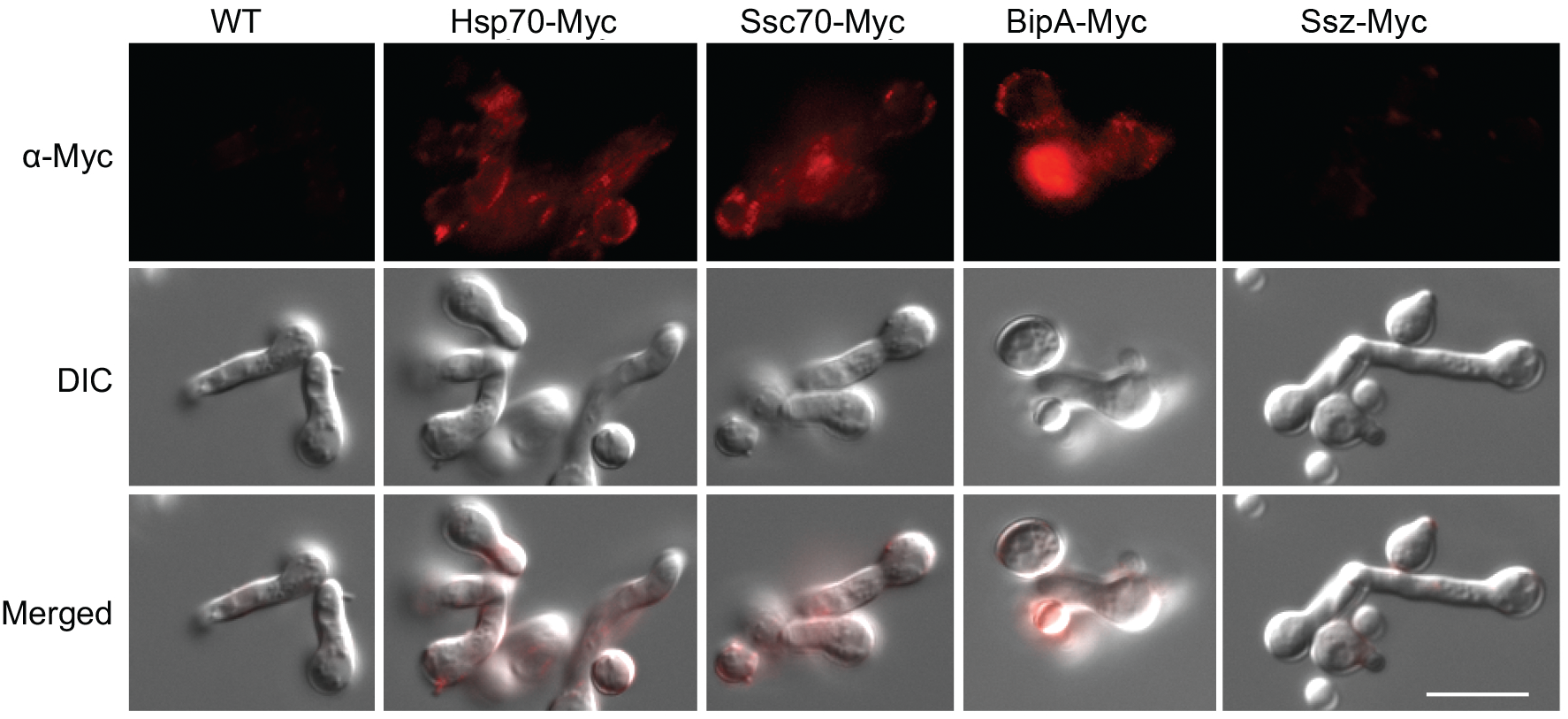
Immunofluorescence staining of *A. fumigatus* germlings. Germlings were blocked in PBS with 1% (w/v) BSA, and then incubated with the primary anti-Myc antibody. The surface Myc-tagged Hsp70s were indirectly detected with an AF568 conjugated secondary antibody. Scale bar = 10 μm.

### 3.5 The core surface proteome (surfome) of *A. fumigatus*

Considering that several surface proteomic studies have been performed using different methods (trypsin-shaving, HF/pyridine, and formic acid extraction), we attempted to compare these datasets to provide a more comprehensive picture of the *A. fumigatus* surfome [9–11]. In total, 992 different proteins were detected as surface proteins in this study and the aforementioned studies, including 437 proteins that were detected by at least two approaches (Figure 5 and Table S3). 43 proteins were commonly identified in all studies (Figure 5 and Table S3). These 43 proteins include 15 of the most prevalent proteins identified in our study and 9 allergens (Table 2, 3, and S3). Two small proteins, Grg1 (glucose-repressible gene) and ConJ (conidiation-specific protein 10), were both detected among the most prevalent proteins on dormant conidia of the ATCC 46645 strain using the HF-pyridine extraction method [10], on the CEA10 strain using the trypsin shaving method [11], and in this study using biotinylation (Table 2). Grg1 was detected with biotinylation at the positions K32 and K46, while ConJ was biotinylated at the N-terminus and site K48 (Table 2). The conidial surface protein CcpA, which contributes to fungal virulence, was detected with biotinylations at positions K41 and K90 (Table S3). The putative GPI-anchored cell wall protein CweA [11] was detected with a biotinylation marker at site K358 (Table S3). In addition to the 43 commonly detected proteins, 68 proteins were detected in at least four surfome datasets (Figure 5A), 44 of which were detected with biotinylated amino-groups (Table S3), including five allergens (Asp f 1, Asp f 4, Asp f 11, Asp f 12, and Asp f 18) and seven other prevalent proteins (1,3-beta-glucanosyltransferase Bgt1, cell wall protein Ecm33, nucleoside kinase Ndk1, eukaryotic initiation factor 5A [Afu1g04070], a putative ADP/ATP carrier [Afu1g05390], the 60S ribosomal protein L13 [Afu4g04460], and ubiquitin [Afu3g11260]) (Table 2). In total 133 surface proteins with biotinylation modifications were also confirmed by alternative methods (Figure 5B). All in all, the core-surfome of *A. fumigatus* builds a valuable database to provide protein targets for further studies on host-pathogen interactions, diagnosis, and immunotherapy.

**Figure 5.**
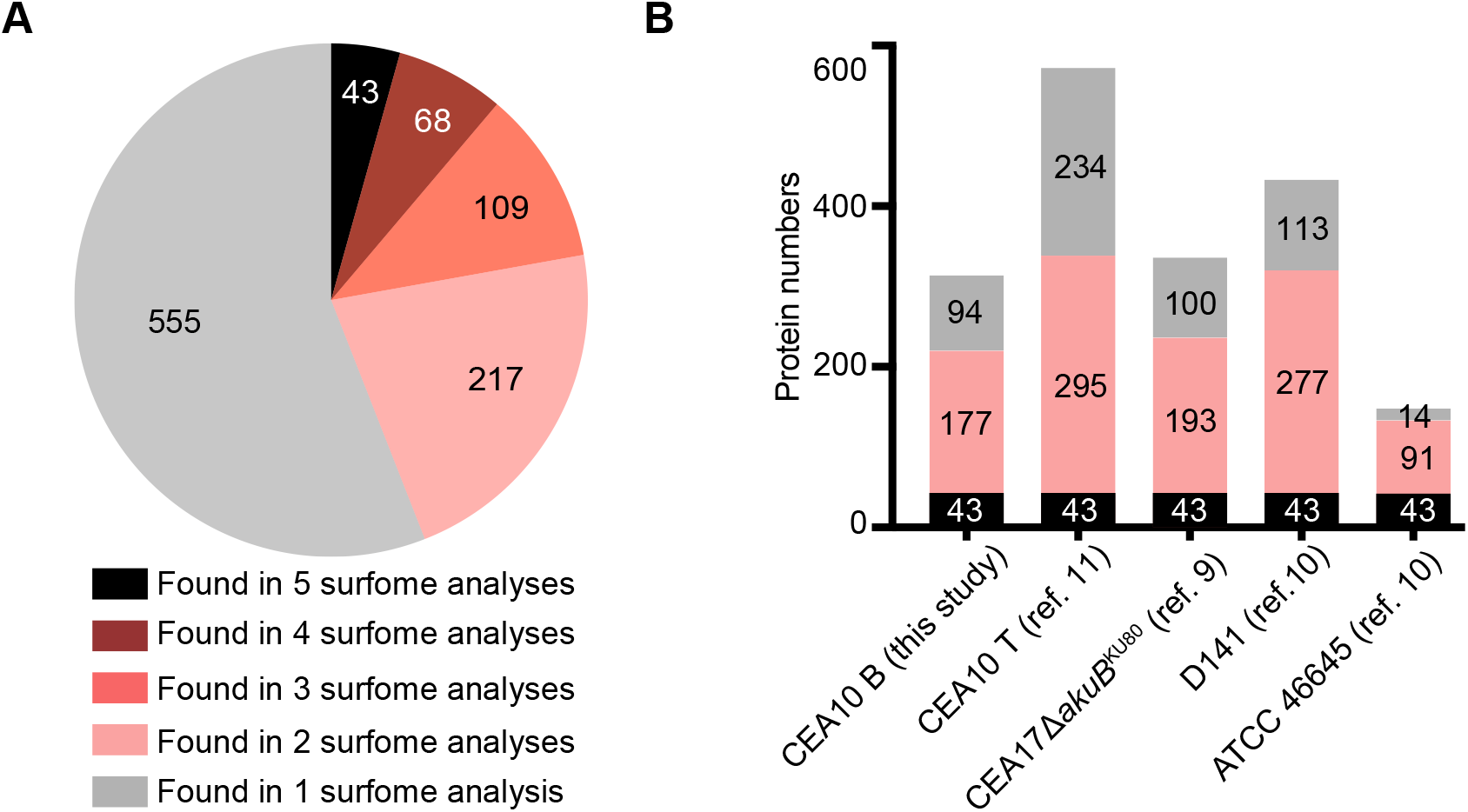
The number of *A. fumigatus* surface proteins identified in different surface proteomics experiments. (A) Pie chart shows the number of proteins present in different experiments. (B) Each of the bars represents the number of identified proteins, with black zones highlighting the number of proteins present in all of the surfome data available, including the results obtained in this study (CEA10 B, CEA10 using biotinylation method), the surfome of CEA10 (CEA10 T, CEA10 using trypsin shaving method) and D141 obtained using trypsin shaving method [10, 11], the CEA17Δ*akuB*^KU80^ dormant conidia surfome obtained from the formic acid extract [9], and the ATCC 46645 surfome detected by hydrogen fluoride-pyridine extraction and trypsin shaving [10]. The pink zones highlight the number of proteins present in at least two but not all the surfome data. Grey zones indicate the number of proteins present in just one experiment.

## 4. DISCUSSION

Proteins in combination with other cell wall components on the surface of *A. fumigatus* conidia play a key role in protecting the fungus from environmental insults and host defense responses during an infection [10, 12, 15, 40, 41]. Several methods and techniques have been used to investigate the surface proteins of this pathogenic fungus, leading to the detection of hundreds of surface proteins, whose presence on the surface changes dynamically during development [9–11, 19, 42]. Most methods used for the extraction of surface proteins, like enzymatic treatment with glucanase or trypsin or acidic extraction with formic acid or hydrogen-fluoride–pyridine, have the potential to release intracellular, unexposed cell wall proteins. Cell impermeable biotinylation reagents, which react with primary amines accessible on the cell surface, have successfully been used for labeling and identification of surface proteins with little contamination [15, 21, 22, 43, 44]. To further expand the *A. fumigatus* surface proteome and to solidify the data of the common core surfome of *A. fumigatus*, we used a surface biotinylation approach, which has not been applied to *A. fumigatus* previously. The aim was to characterize and detail the *A. fumigatus* surfome with surface exposed regions across germination and verify the surface localization of selected proteins by an additional method (Figure 4). Therefore, our work provides a multitude of candidates for further investigation of host-pathogen interaction and possible immuno-diagnostic/therapeutic usages.

Proteins on the surface mediate the direct contacts between pathogens and hosts. In addition to several known surface proteins, like RodA, CalA, Asp f 2, and CcpA [10, 12, 15, 18], abundant surface proteins detected in this study, such as Tef1, ArtA, Hsp70 and Hsp90, also have the potential to interact with a range of host proteins [45]. The *Candida albicans* Tef1 was shown to be able to bind human plasminogen, probably through the C-terminal lysine residues [46]. In this study, Tef1 was detected with biotinylation at the C-terminal K472–K486 region throughout germination (Table 1), indicating that Tef1 in *A. fumigatus* might have a similar function. The 14-3-3 proteins, such as ArtA, also have the ability to bind a multitude of proteins, and play an important role in morphogenesis and sensing of environmental cues in fungi [47–49]. The surface-localized heat shock proteins influence the interaction between bacterial and fungal pathogens and host cells as well [50–52]. For example, expression of *Listeria monocytogenes* heat shock protein ClpC, a member of the 100-kDa heat shock protein family, is required for cell adhesion and invasion [53], and also allows this bacterium to escape from the phagosome [54]. The *C. albicans* Hsp70 protein Ssa1 is an invasin that binds to host cell cadherins to induce host cell endocytosis, which is critical for *C. albicans* to cause maximal damage [55]. In line with this, the *A. fumigatus* Hsp70 and Hsp90 were predicted to interact with host proteins in conidia containing mouse macrophage phagolysosomes [45], suggesting the potential roles of surface heat shock proteins in manipulating the host immune responses.

Besides their roles in pathogenicity, surface proteins could also be targets of immunotherapies based on antibodies. Surface proteins are particularly suitable due to their easy accessibility. Antibody-based therapies are rapidly growing, since they are a promising approach to directly attack the pathogen and boost the innate immune system. However, only a few monoclonal antibodies against fungal pathogens have been developed and advanced to clinical trials [56]. One example is Mycograb, an Hsp90-specific antibody fragment, which showed promise for treating *Candida* infections in combination with amphotericin B [57], but failed to obtain marketing authorization. In mouse experiments, treatment with anti-HSP 60 antibodies reduced fungal burden after infection with the dimorphic fungi *Histoplasma capsulatum* and *Paracoccidioides lutzii* [58, 59]. On the negative side, the amino acid sequences of HSPs are highly conserved and cross-reactivity may occur. For example, the epitope (NKILKVIRKNIVKK) that Mycograb targets shows high similarity between yeast, mice, and human homologues of Hsp90 [57], including *A. fumigatus* Hsp90 (NKIMKVIKKNIVKK, amino acids 383–396). Other, abundant, fungal-specific surface proteins may represent better targets for immunotherapy as discussed in the following.

To date, more than 100 antigens or immune-reactive proteins of *A. fumigatus* have been identified using classical immunobiological procedures that react with sera from ABPA patients or animal models [35–37]. However, only a few recombinant allergens have been used commercially for diagnosis of allergic aspergillosis [5]. Although the crystal structures of some allergens are known [60, 61], the question of whether the allergen/protein exhibits special structural characteristics that are responsible for its allergenicity is still poorly understood. Thus, the information about the association of allergens with the different morphotypes (conidia, mycelium) and their exposed regions that may directly mediate the interaction with host components, such as IgE binding, is needed. Such knowledge creates the basis for the understanding of the immunological properties of protein antigens and is important for the establishment of new forms of diagnosis and treatment [62].

The cyclophilins, including Asp f 11 and Asp f 27 [61], which are structurally conserved pan-allergens showing extensive cross-reactivity [63, 64]. It was reported that the conserved N81-N149 region of Rhi o 2 from *Rhizopus oryzae* (Figure S4) is crucial for IgE-recognition and cross-reactivity [63]. In this study, however, the C-terminal region of Asp f 27, which is not conserved, was detected with biotinylation marks instead (Table 3 and Figure S4). Several other allergens were detected with biotinylation sites on different morphotypes, such as the K50–K101 region of Asp f 6, K304–K327 region of Asp f FDH, K238–K269, and K303–K331 region of Asp f MDH (Table 3). Noteworthy, there were also regions of surface proteins that were detected on all morphotypes. For example, the K50–K55 region of RodA, K472–K486 region of Tef1, K122 region of Afu5g10550, K125–K141 region of Bgt1, K24–K63 region of Afu3g00880, K225–K231 region of Bgt2, and K420–K435 region of the putative dihydrolipoamide dehydrogenase Afu2g02100 were all consistently surface exposed (Table 1). These peptides may represent promising candidate antigens for the development of monoclonal antibodies to be used for diagnosis and immunotherapy.

As published in our previous study, the surfome of *A. fumigatus* is dynamic [11]. Correspondingly, a recent study also revealed that the phenotypes of *A. fumigatus* germinating conidia vary among genetically identical conidia. Here, we detected 74, 75, and 93 proteins on dormant, swollen, and germinating conidia, respectively; while 214 proteins were found on the surface of hyphae (Figure 3). Comparison of the different surface proteomics datasets helped to identify the core-surfome of *A. fumigatus* (Figure 5). These proteins, which are consistently found on the surface of *A. fumigatus*, likely play a role in mediating the interaction of *A. fumigatus* with the host or other organisms.

## 5. CONCLUSION

A surface biotinylation enrichment coupled with LC-MS/MS analysis was applied to study the surface proteome of *A. fumigatus* across germination. 314 surface proteins, including RodA, some virulence factors, allergens, and newly identified surface proteins were detected with biotinylation modifications, confirming exposed surfaces for these proteins. These data highlight not only the surface proteins but also the exposed regions that could be used in the future for new diagnostic tools and immunotherapies based on monoclonal antibodies or T cell-based vaccines.

## ASSOCIATED CONTENT

### Supporting Information

**Table S1.**
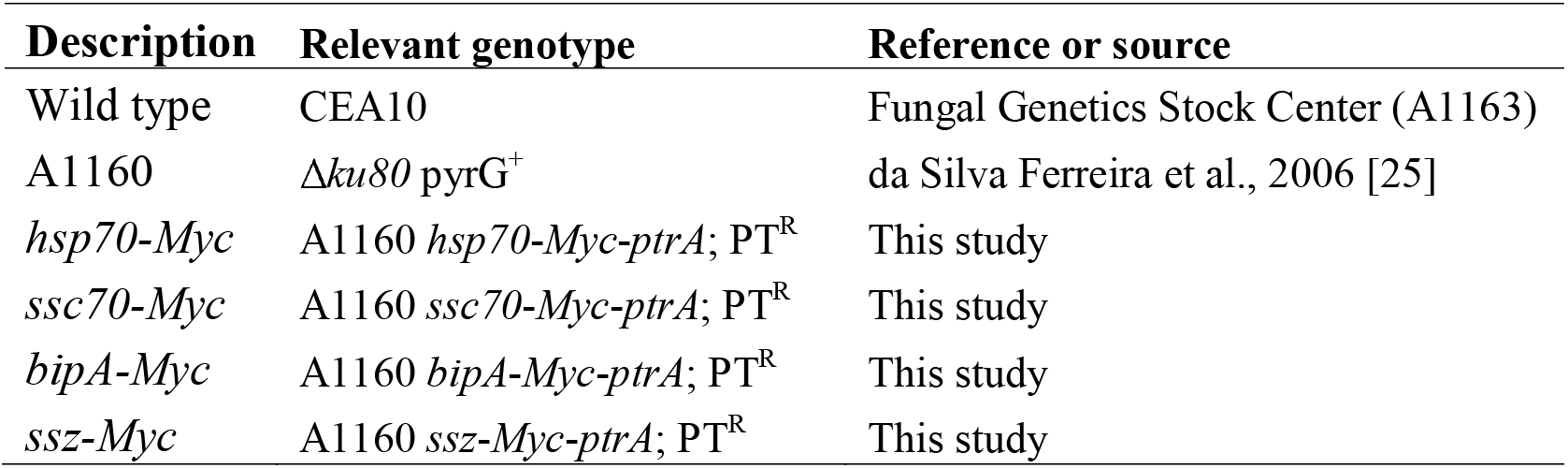
*A. fumigatus* strains used in this study.

**Table S2.**
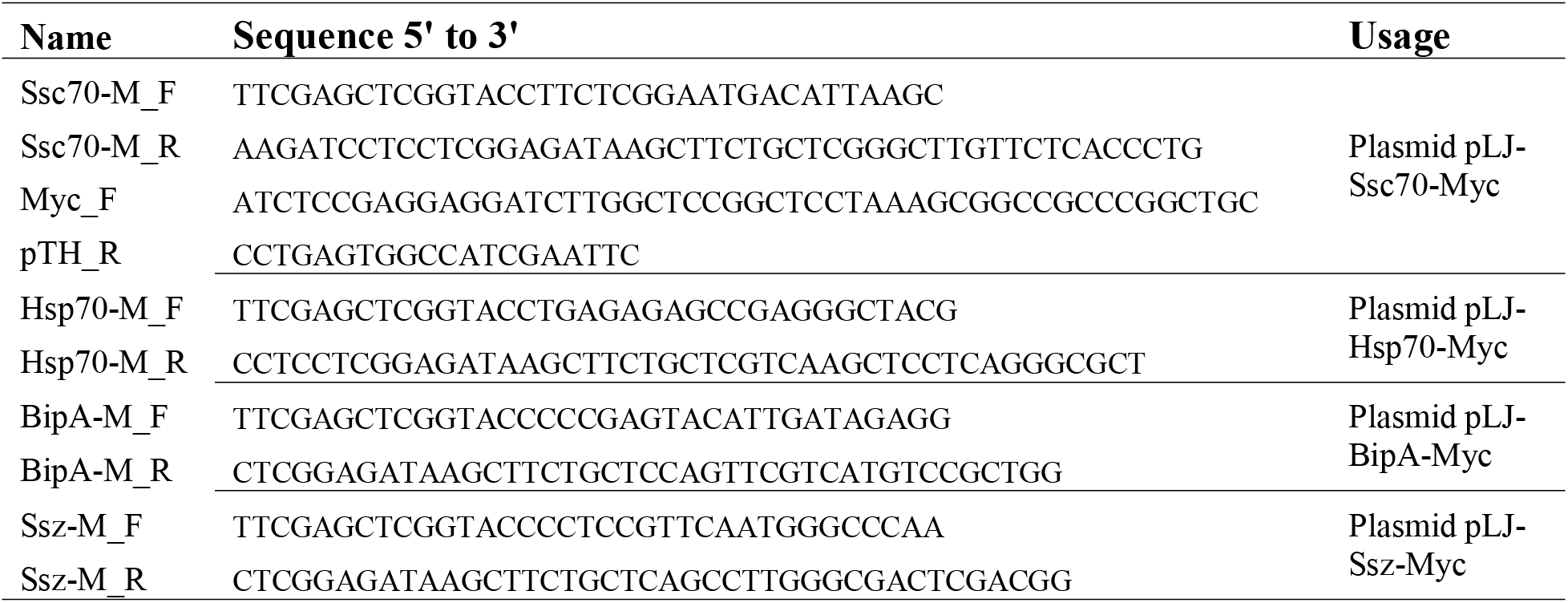
Oligonucleotides used in this study.

Table S3. Proteins of *A. fumigatus* detected with biotinylation marks.

Dataset 1. Streptavidin enriched *A. fumigatus* proteins detected by LC-MS/MS. For the headers, B1, B2, B3, and B4 are different repeats of biotinylated samples; C1, C2, C3, and C4 are different repeats of control samples.

**Figure S1.**
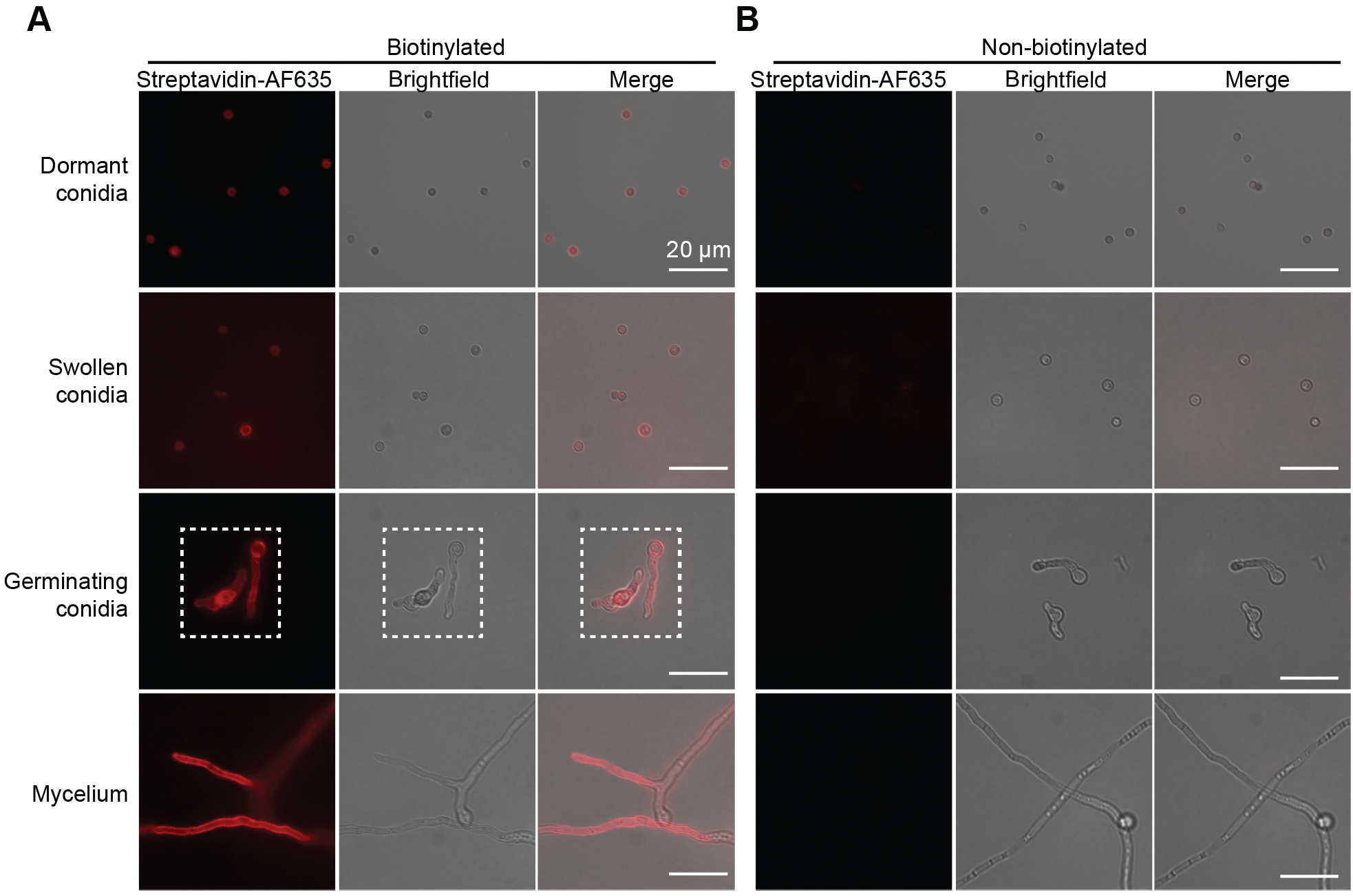
Immunofluorescence staining of biotinylated (A) and non-biotinylated (B) *A. fumigatus* morphotypes with Alexa Fluor^™^ 635 conjugated streptavidin (Streptavidin-AF635). Germlings indicated with dashed boxes are also shown in Figure 2A. Scale bar is 10 μm.

**Figure S2.**
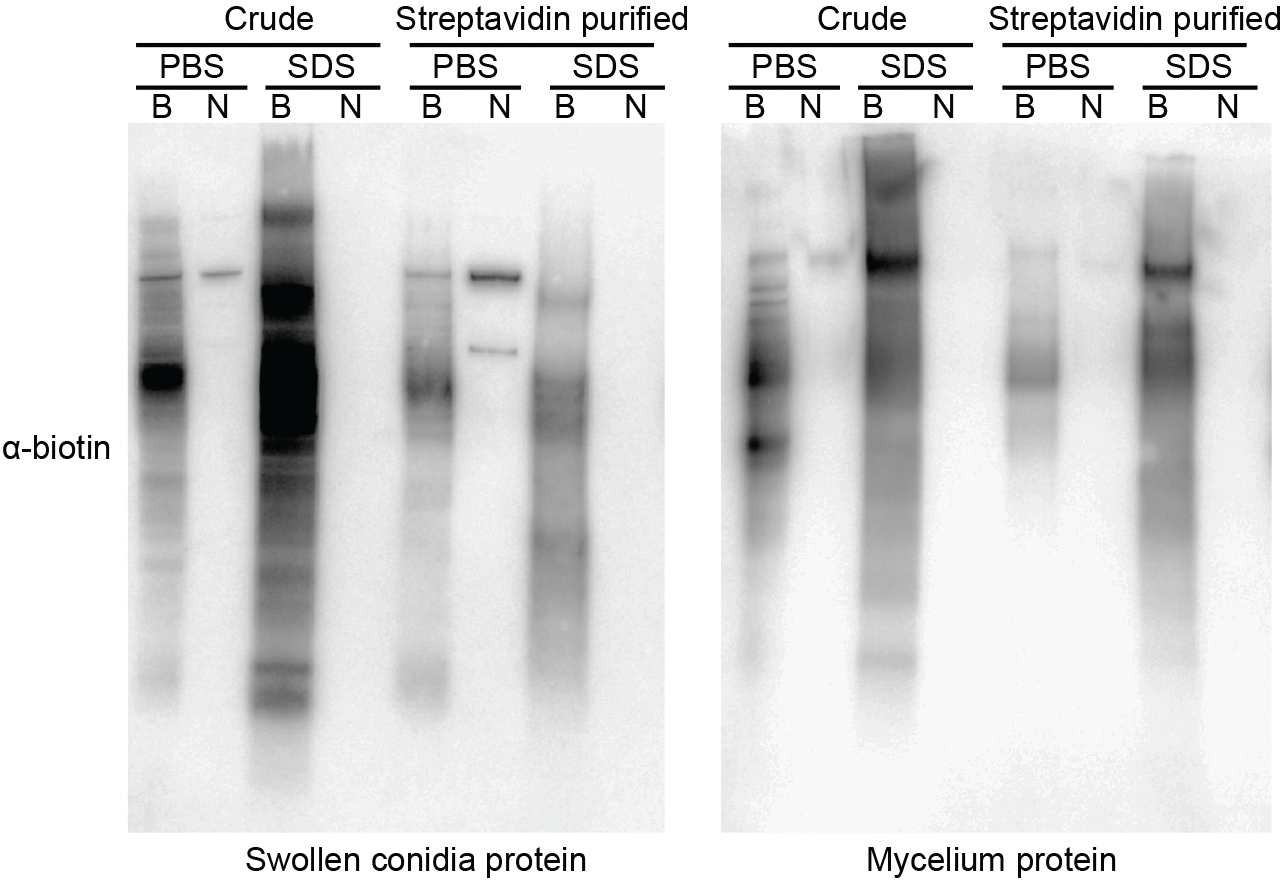
Immunoblotting analysis of swollen conidia and mycelia protein extracts. Crude protein extracts (crude) and purified proteins (streptavidin purified) from PBS buffer (PBS) and SDS buffer (SDS) were separated on SDS-PAGE and then transferred to PVDF membrane. Biotinylated proteins were detected with Pierce^®^ streptavidin-HRP. B, biotinylated; N, non-biotinylated.

**Figure S3.**
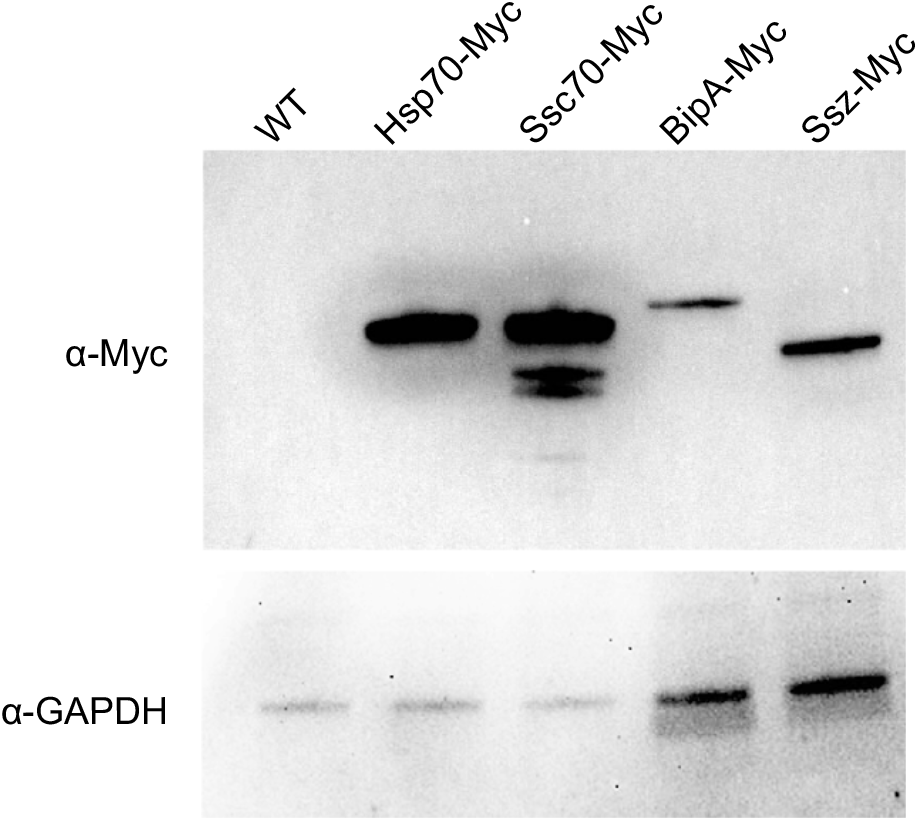
Immunoblotting of *A. fumigatus* protein extracts. Myc-tagged proteins were detected using α-Myc antibody. For the *hsp70-Myc* and *ssc70-Myc* expressing *A. fumigatus* strains, protein was extracted from dormant conidia; while for the *ssz-Myc* and *bipA-Myc* expressing strains, protein was extracted from mycelium. An α-GAPDH antibody served as loading control.

**Figure S4.**
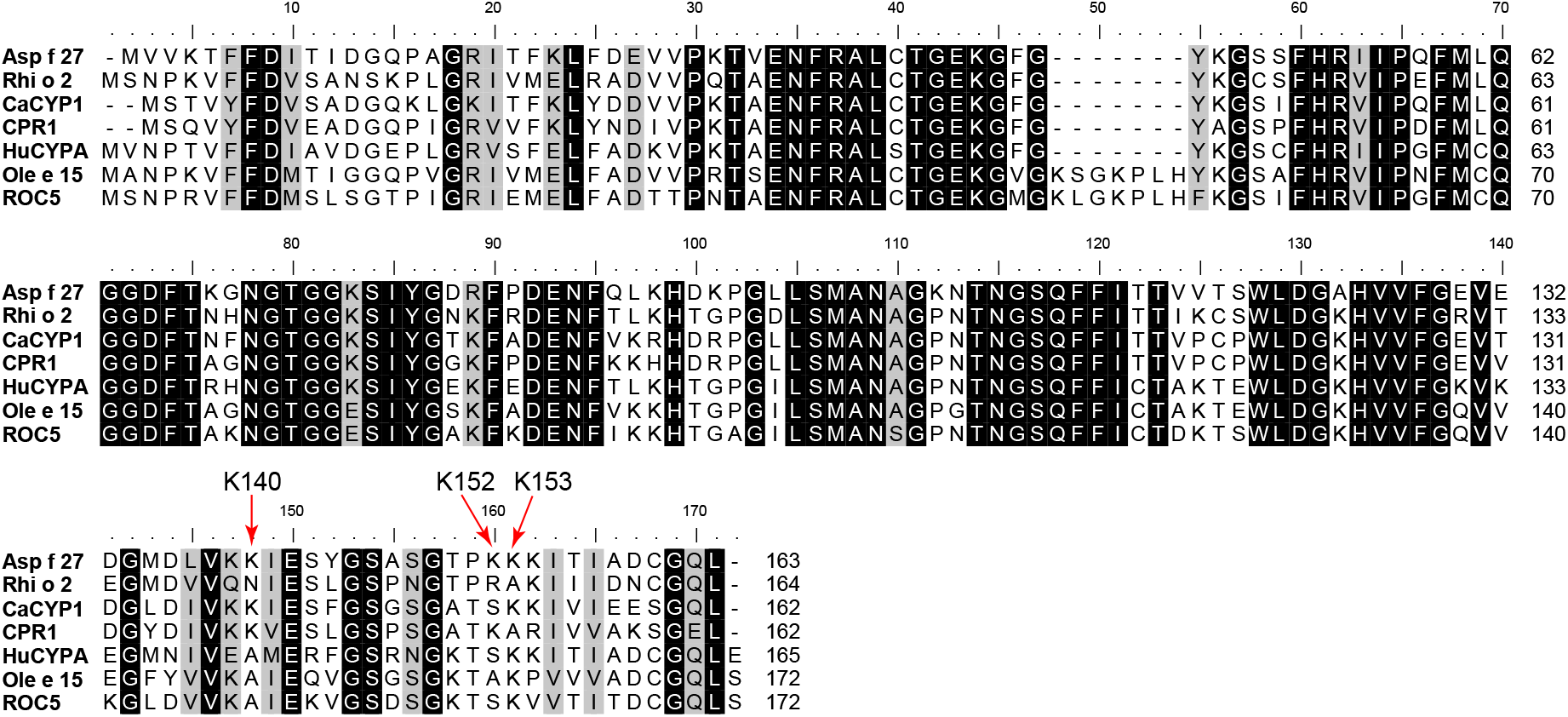
Alignment of Asp f 27 and homologous proteins. Lysine residues of Asp f 27 detected with biotinylation marks in this study are indicated with arrows. Protein sequences of Asp f 27 (UniProtKB: Q4WWX5), CaCYP1 (*Candida albicans*, UniProtKB: P22011), and CPR1 (*Saccharomyces cerevisiae*, UniProtKB: P14832), and Rhi o 2 (*Rhizopus delemar*, UniProtKB: P0C1H7) are downloaded from UniProt Knowledgebase (www.uniprot.org). Protein sequences of Ole e 15 (*Olea europaea*, GenBank: AVV30163.1), HuCYPA (Homo sapiens, GenBank: AAH05982.1), and ROC5 (*Arabidopsis thaliana*, NCBI Reference Sequence: NP_195213.1) were downloaded from NCBI (www.ncbi.nlm.nih.gov).

## Notes

The authors declare no conflicts of interest.

## ACKNOWLEDGMENT

This work was supported by the Deutsche Forschungsgemeinschaft (DFG)-funded French-German project “AfuInf”, the DFG Collaborative Research Center/Transregio FungiNet 124 ‘Pathogenic fungi and their human host: Networks of Interaction’ (project A1, Z2), and the Federal Ministry of Education and Research, project EXASENS (13N13861). The funder had no role in the design or choice to publish these data.

